# Bacteriophage-Loaded Microneedle Patches for Targeted and Minimally Disruptive Foodborne Pathogen Decontamination

**DOI:** 10.1101/2025.04.30.651002

**Authors:** Akansha Prasad, Shadman Khan, Fatima Arshad, Hareet Sidhu, Kyle Jackson, Roderick MacLachlan, Ekaterina Kvitka, Veronica Grignano, Hannah Mann, Carlos D. M. Filipe, Zeinab Hosseinidoust, Tohid F. Didar

## Abstract

Antibacterial additive use has surged due to rising incidences of food contamination, despite concerns over antibiotic resistance. Bacteriophage (bacterial viruses) represent a unique and promising opportunity as antibacterial agents, offering targeted bacterial lysis while being food safe. However, their commercial success has been limited by the significant diffusion barriers they face within food, preventing effective delivery at contamination sites. Here, we introduce bacteriophage-loaded microneedle patches that enable targeted phage delivery directly within food, eliminating internal pathogens in a minimally disruptive manner. The application of microneedles within food is first explored. The platform is then substantiated by comparing performance in raw beef and cooked chicken, where we achieved up to 3-logs reduction in *Escherichia coli*, thus providing complete decontamination according to regulatory limits. In contrast, conventional surface application of the same phage failed to provide significant decontamination. To ensure broad applicability, phage cocktails were also loaded into microneedles to demonstrate polymicrobial decontamination against other common food contaminants including *Salmonella*. This platform can also be adapted to extend food shelf-life by targeting spoilage-inducing bacteria.

---

Despite recent technological advancements and more stringent regulatory measures, foodborne illness persists as a major global crisis, resulting in over 600 million annual cases and causing 7.5% of all annual deaths.^1,2^ The presence of pathogenic contaminants, such as *Escherichia coli* O157:H7 and *Salmonella enterica*, along with the costly and retroactive food recalls that they trigger, are both projected to rise as our reliance on intricate, multi-faceted global supply chains continues to grow.^3–8^ Nearly half of all food recalls in the United States are attributed to the presence of microbial contamination, and ready-to-eat (RTE) products are amongst the most recalled items, making them of particular interest.^9^

Antibiotics have traditionally been used as commercial antibacterial agents during agriculture, however, their use has faced recent scrutiny based on growing concerns surrounding the rise of antibiotic-resistant bacteria.^10,11^ Furthermore, many food products, particularly RTE items, are highly susceptible to contamination during later stages of the food production process, necessitating the incorporation of antibacterial agents directly into the food product. To this end, the use of antibacterial agents such as metal additives, chemical compounds and essential oils has garnered significant research interest.^12,13^ These agents are scalable, food-safe, and exhibit high bactericidal activity.^14–16^ However, their commercial viability is limited by either their poor specificity or stability, as well as their disruptive effects on the product’s organoleptic properties.^17–19^

Bacteriophages, on the other hand, have proven effective in eliminating bacterial contamination in various food products.^20–27^ Bacteriophages (or phages for short) are viruses that target bacteria in a highly specific manner. This specificity means phage antimicrobials can be designed to eliminate contamination without affecting any non-pathogenic microbes that are innately present within the food and are responsible for palatability.^28–31^ Bacteriophages have regulatory approval for food applications and importantly do not affect food taste, texture, or odor.^32,33^ Adding to the long list of desirable characteristics, phages propagate when contacted with bacterial hosts, and are thus self-dosing antimicrobial additives.^28^ Phage is commonly applied to solid food surfaces through methods such as spraying and liquid emersion.^30,34^ However, phages are proteinaceous nanoparticles, ranging in size from tens to hundreds of nanometers, and thus experience significantly larger diffusion barriers in solid matrices compared to small molecular antimicrobials. Therefore, the main bottleneck for achieving phage antimicrobial efficacy in solid food matrices is overcoming these barriers by delivering phage to the site of contamination.^35,36^

Microneedles represent a robust tool to achieve this bacteriophage delivery directly into food matrices, offering penetrative ability at a nondisruptive scale, all in a low-cost form factor that is easily applied by hand. Microneedles have been extensively explored in clinical applications as an attractive, minimally invasive solution for crossing the skin barrier. Yet, they remain relatively unexplored for on-food applications, despite evidence from biomedical literature that suggests compatibility with fluid rich, solid food matrices.^37–40^

We have developed food-safe, bacteriophage-loaded microneedles, which enable phage delivery directly into food matrices for effective antibacterial activity. We performed extensive microneedle characterization both mechanically and with various food products to select an optimal polymer. We then demonstrate bacteriophage-induced antibacterial activity within raw and RTE food products with the loaded microneedle patches compared to other phage delivery methods. We further show that our microneedle patches are capable of polymicrobial decontamination for both *Escherichia coli* and *Salmonella enterica* serovar Typhimurium, when loaded with bacteriophages specific to both pathogens.

## Results and Discussion

### Bacteriophage-loaded microneedles design

We aimed to design a bacteriophage-loaded, food-scale microneedle platform that provides antibacterial activity in various food products (**Figure 1a**), especially in high-risk RTE products. In the presence of their pathogenic targets, corresponding bacteriophages immobilized along the polymer microneedle surface infect adjacent targets, enabling species level specificity (**Figure 1b**). The platform was validated both in raw beef (**Figure 1c**) and with RTE chicken, where the bacteriophage-loaded microneedles outperformed flat, surface-level patches, thus overcoming the diffusion barrier (**Figure 1d**). Further applications of the microneedles for polymicrobial decontamination (**Figure 1e**) and as large-scale arrays were also explored (**Figure 1f**). To ensure food compatibility and viability, four different polymeric materials and two different, highly effective bacteriophages were used in this work. Notably, all these materials were deemed food safe by the FDA as they were designated Generally Recognized As Safe (GRAS) and placed on the Indirect Food Substances regulation.^3^

**Figure 1.**
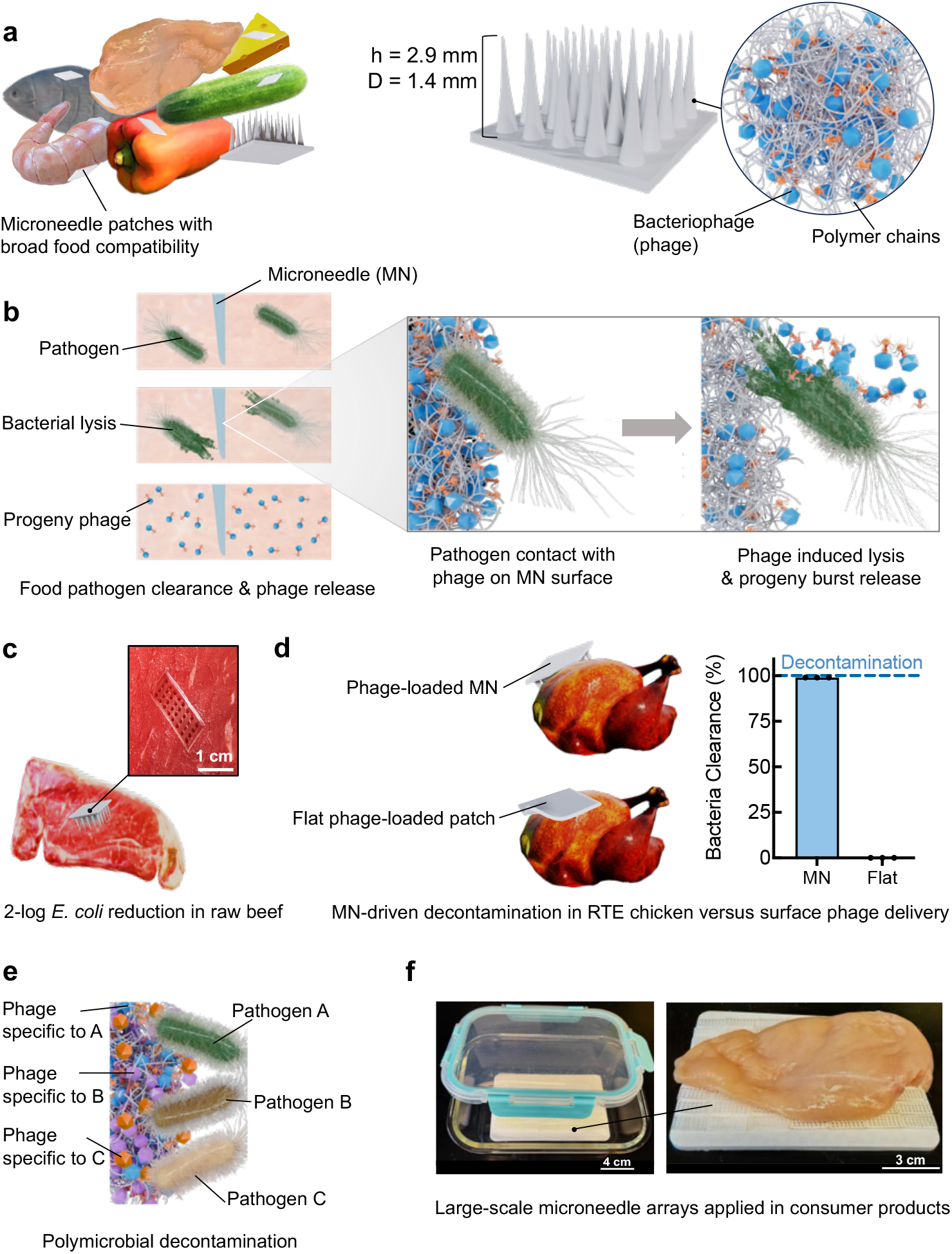
Bacteriophage-loaded microneedles design. (a) PMMA polymeric chains with bacteriophage incorporation applied to assorted food items as antibacterial additives. This microneedle platform has a relatively larger scale compared to clinical microneedles to ensure food-compatibility. (b) Macroscale overview of bacterial clearance and decontamination within food matrices. Microscale view is magnified to show selective bacteriophage delivery upon target presence, resulting in contaminant lysis and subsequent phage propagation. (c) Microneedle applied to raw beef with magnified optical image shown. (d) Penetrative microneedle performance assessment compared to flat patches in RTE chicken, where phage loaded microneedles achieve complete decontamination. (e) Microneedles with polymicrobial decontamination potential when loaded with multiple phages specific to different food pathogens. (f) Multiple microneedle arrays unified to create a large-scale platform that provides continuous, whole-product decontamination.

### Microneedle material mechanical characterization

A detailed comparison of microneedle material candidates was first performed to understand the effect of material choice on microneedle properties. The biocompatible polymers polymethyl methacrylate (PMMA), polyvinyl alcohol (PVA), polydimethylsiloxane (PDMS), and gelatin were selected (**Note S1, Table S1, Figure S1**). Micromolding with a custom stereolithography (SLA)-printed mold (**Figure 2a, Figure S2**) was used to fabricate scalable microneedle arrays of the candidate materials (**Figure 2b**). Scanning electron microscopy (SEM) revealed that gelatin exhibited a unique, highly textured surface due to the amorphous nature of its polymeric chains, while PMMA, PVA, and PDMS appeared relatively similar in surface texture and structure (**Figure 2c**).^41^ The potential use of this platform for fluorescence monitoring was also supported by all materials displaying low fluorescence across the visible light spectrum (**Figure S3**).

**Figure 2.**
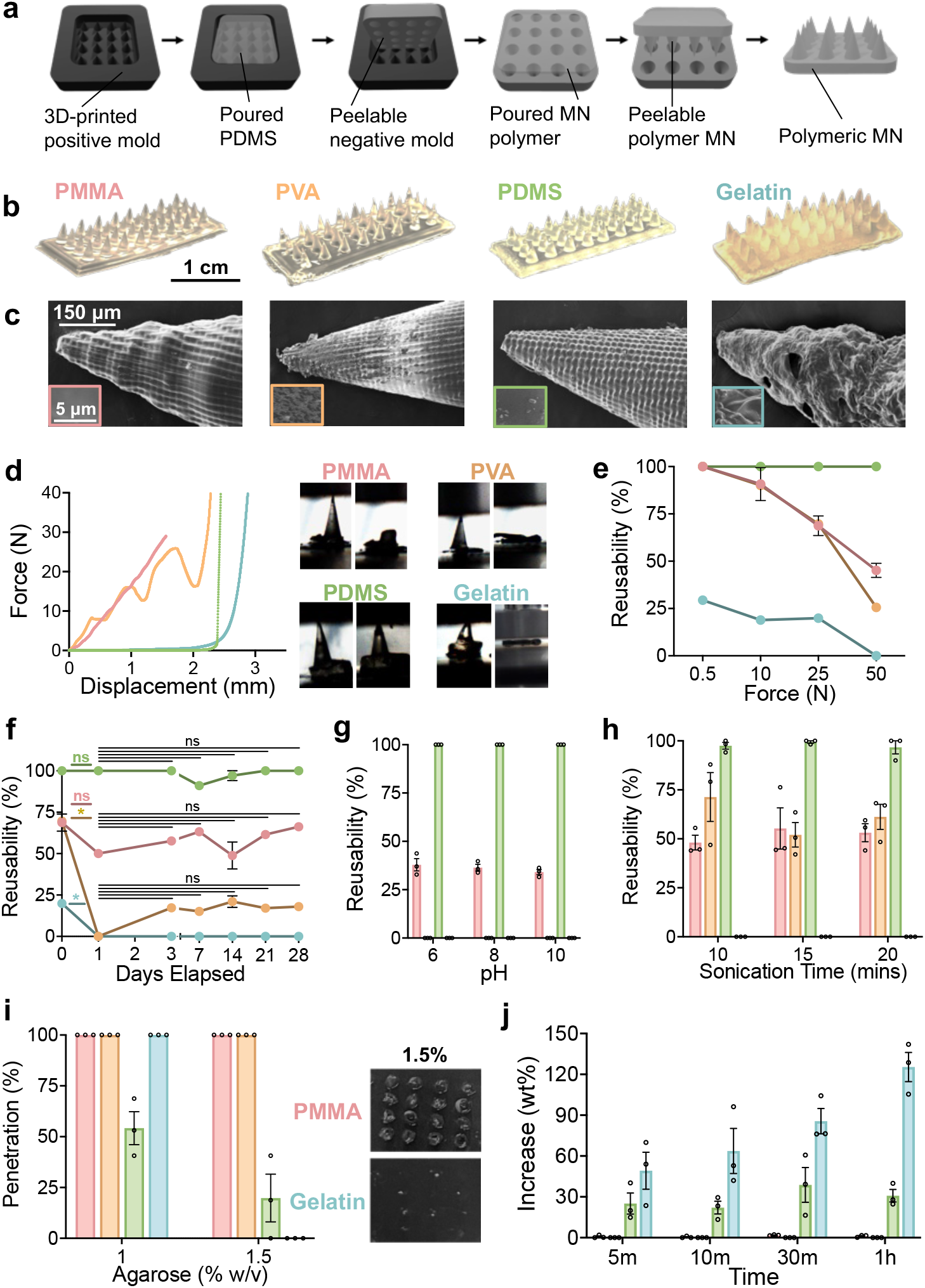
Microneedle mechanical characterization and material comparison. (a) Overview of microneedle fabrication via micromolding. (b) Optical images of microneedle array patches. (c) SEM images of microneedles at 350X with 10000X overlays. (d) Force-displacement curves of all material candidates under 50N of compressive force with associated pre- and post-compression microneedle images. (e) Microneedle reusability across increasing compressive forces for all candidate materials. (f) Short and long-term stability of all microneedle materials with asterisk representing significant differences at corresponding significance level. (g-h) Continued microneedle stability testing through exposure to (g) pH levels and (h) sonication-based stressors. (i) Preliminary penetration assessment of all microneedle material candidates based on penetrative performance in agarose disks of varying density with pictures showing penetration in 1.5% agarose. (j) Water absorbance of PMMA, PDMS, and gelatin overtime. All reported values represent the mean of all samples with error bars representing standard error of the mean.

Compression testing highlighted differences in mechanical properties, with PMMA and PVA showing steep linear deformation curves, indicative of their high tensile strength (**Figure 2d, Figure S4**).^42,43^ Both PDMS and gelatin demonstrated elastic behaviour, with PDMS demonstrating superior recovery due to its low elastic modulus.^44,45^ This elasticity enabled PDMS to display the highest reusability under compressive loads, quantified as the percentage of microneedles within a patch that remained undamaged and thus could be reused (**Figure 2e**). Stability testing involved storing the microneedle samples for short and long-term periods at room temperature and repeating compression testing (**Figure 2f**). Both PDMS and PMMA had no significant deficits in their strength (*P* > 0.05), while PVA and gelatin exhibited significant depletion over an initial 24 hour period (*P* < 0.05). Temperature effects were also explored across a range of temperatures, with similar trends (**Figure S5**). The poor stability of PVA and gelatin is attributed to their aqueous hydrogel nature, which deteriorates with long-term storage, yielding a depletion in mechanical strength.^46^ A similar test was performed after 24 hour storage in environments with pH levels of 6, 8, and 10 to assess resiliency in acidic and basic environments (**Figure 2g**). PDMS had no deterioration in performance, while PMMA exhibited ~ 50% reduction in reusability. Resilience against mechanical stress *via* exposure to sonication was also explored, with PDMS and PMMA displaying similar reusability compared to their baseline values after 10, 15, and 20 minutes (**Figure 2h**). Intriguingly, the PVA microneedles were also able to retain their original performance despite the mechanical stress, which is attributed to the strong hydrogen bonds innately present within the hydrogel. The gelatin microneedles were fully degraded, given their poor mechanical strength.

Penetrative strength was assessed using agarose discs of varying strength, where PMMA and PVA excelled with 100% penetration, given their high tensile strength (**Figure 2i**). The gelatin microneedles were also able to provide complete penetration of the softer 1% agarose disk, but broke when attempting to penetrate through a 1.5% disk. Despite their high reusability, the PDMS microneedles had extremely poor penetration due to their innate elasticity, which provided insufficient penetrative strength. The absorption capacity of the various microneedle candidates was also explored, where the gelatin microneedles absorbed significantly higher amounts of water given their hygroscopic nature (**Figure 2j**).^41^ The permeability of the microneedles was also explored for their potential in dissolvable or sensing-based platforms (**Method S1, Figure S6**). The mechanical characterization results of all four material candidates are summarized in **Table S2**.

### Comparison of microneedle materials for food-specific applications

While the PMMA microneedles demonstrated the best mechanical properties, all four materials were evaluated for food-specific applications as each material offered distinct strengths and weaknesses that could influence their effectiveness across foods of varying textures. We selected five solid, moisture-rich food products – mushrooms, peaches, fish, RTE chicken, and cheese – for food-specific testing (**Figure 3a**). The selected range of products represents a spectrum of varying porosity and density, ranging from soft and highly porous (mushrooms and peaches) to dense and firm (RTE chicken and cheese) to ensure robust characterization.^47,48^ Penetration and reusability studies (**Figure 3b,c**) were conducted at insertion forces of 2.5, 5, and 7.5 N, a range in line with those used in clinical studies to consistently approximate user-applied force.^37,49,50^ Reusability is of particular concern when creating products for the food industry to ensure cost efficiency. An ideal material candidate for bacteriophage delivery should thus offer both high penetration and reusability. PMMA demonstrated regularly higher 100% penetration and reusability for all food items, while PVA also demonstrated high penetration across almost all foods, with mixed reusability. Similar to the previous characterization studies, PDMS exhibited high reusability, but poor and inconsistent penetration due to its elasticity. Interestingly, gelatin had improved penetration in fluid-rich foods such as peach and fish as well as in softer foods like cheese. However, it had relatively low and very inconsistent reusability. To further elaborate on reusability, the PMMA microneedle patches remained perfectly intact, only the microneedle tips were damaged in PVA and PDMS needles, and the gelatin microneedle patches were destroyed across most food items (**Figure 3d**). These tests indicated that PMMA offered high performance in both penetration and reusability assessments (**Figure 3e**).

**Figure 3.**
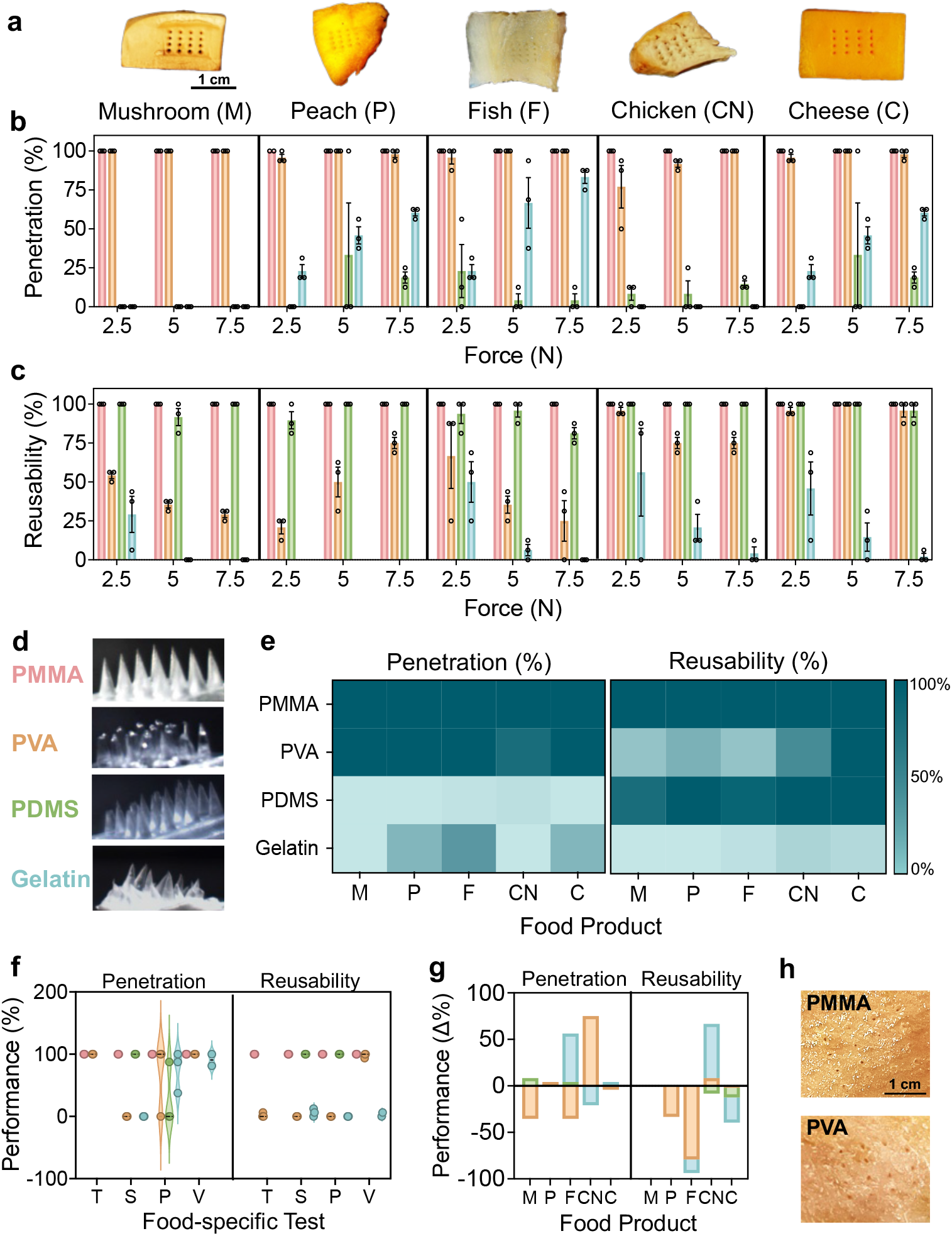
Food-specific microneedle characterization. (a) Optical images of various food items post penetration from PMMA microneedle arrays. (b-c) Microneedle material evaluation through (b) penetration and (c) reusability assessment at 2.5, 5, and 7.5 N of force. (d) Associated optical images of microneedle reusability after insertion into peach samples at 5 N. (e) Heat map summarizing penetration and reusability trends across all microneedle materials for all five food products. (f) Comparison of penetration and reusability through food-centric applications and tests including tumbling (T), emersion in fish saline (S) and chicken purge (P), and being placed under vacuum (V). (g) Change in microneedle performance after 24h incubation in all food items. (h) Associated post-penetration images of porcine skin with PMMA and PVA microneedles. All reported values represent the mean of all samples with error bars representing standard error of the mean.

The validity of these platforms was then substantiated through more food-centric studies, wherein microneedles were inserted into food items and then subjected to various food-relevant stressors (**Figure 3f**). First, microneedle-containing mushrooms and peaches were subjected to mechanical tumbling at similar speeds to those utilized in automated produce processing, where both PMMA and PVA had high penetration but only the former exhibited any reusability. Both PDMS and gelatin patches failed to stay within the food products during tumbling, given their elastic nature. Next, the microneedle samples were exposed to food matrix fluids in the form of fish saline fluid and chicken purge. With respect to immersion in fish saline, both PMMA and PDMS had high penetration. Opposingly, both PVA and gelatin had poor performance despite their previously high penetration in fish. This drastic decrease in performance is attributed to the significantly higher volume of salt they were exposed to in this incubation. The hygroscopic nature of the salt-rich exterior environment is credited to the exasperated hydrogel breakdown and in turn, poor mechanical performance.^51^ PMMA outperformed the other candidates in chicken purge, with all other materials exhibiting inconsistent, moderate to low penetration and mixed reusability. Finally, the microneedles were inserted into cheese samples and placed under vacuum conditions repeatedly to simulate the vacuum sealing process commonly used in food packaging processes by both producers and consumers. Both PMMA and PVA demonstrated excellent performance and gelatin showed high penetration, but poor reusability. PDMS patches were unable to remain penetrated throughout the vacuuming process. A brief stability study was also performed where the microneedles were stored within respective food products for 24h and then assessed through the same penetration and reusability studies (**Figure 3g**). Importantly, both PMMA and PDMS saw no changes to their performance. Owing to their hydrogel composition, both PVA and gelatin typically demonstrated decreases in their overall performance. Finally, microneedle performance was also assessed with porcine skin, an extremely dense product that is often used to characterize clinical microneedles with similar results (**Figure 3h, Method S2, Figure S7**).

Overall, PMMA demonstrated consistently high performance across all material candidates and was consequently selected as the base material for the bacteriophage-loaded microneedle platform.

### Phage-integration into PMMA microneedles and system characterization

Once PMMA was identified as the ideal microneedle material for agent delivery into food, the system was validated with both *E. coli* specific T7 phage and *Salmonella enterica* serovar Typhimurium specific FelixO1 phage (**Figure 4a, Note S2**). Initial validation demonstrated that the phage-loaded microneedles produce lysis zones on bacterial lawns, confirming their antibacterial activity (**Figure 4b**). Subsequent baseline assessments focused on ensuring specific bacteriophage release in bacteria-rich buffer solutions to confirm significant bacteriophage-mediated bacterial clearance (*P* < 0.0001) for both pathogens (**Figure 4c,d**). The system’s compatibility with food matrix fluids was then assessed using plaque assays, which confirmed that mixing liquid bacteriophage suspensions with peach pulp and chicken purge had no significant (*P* > 0.05) adverse effects on bacteriophage release (**Figure S8**). To further validate bacteriophage viability within the microneedles, stress-induced degradation tests were performed by incubating patches under various mechanical and chemical stressors for an hour and then assessing antibacterial performance (**Figure 4e**). Undamaged bacteriophage-loaded microneedles exhibited a significant 6.47-logs bacterial reduction (*P* < 0.0001) compared to unloaded controls. Opposingly, loaded microneedles that were exposed to harsh conditions such as sonication, boiling water, sodium hydroxide, ethanol, or ultraviolet light performed significantly worse, with bacterial log reductions ranging from 2.26 to zero, indicating complete inactivation of phage (*P* < 0.01 – 0.0001). Due to PMMA’s high melting point and stability in harsh chemical environments, any impacts on microneedle integrity were considered negligible.^52^ Thus, this reduced performance was attributed to bacteriophage degradation. A one-week stability study confirmed no depletion in bacteriophage activity following storage at 4°C, ensuring compatibility with food products that require prolonged storage (**Figure S9**).

**Figure 4.**
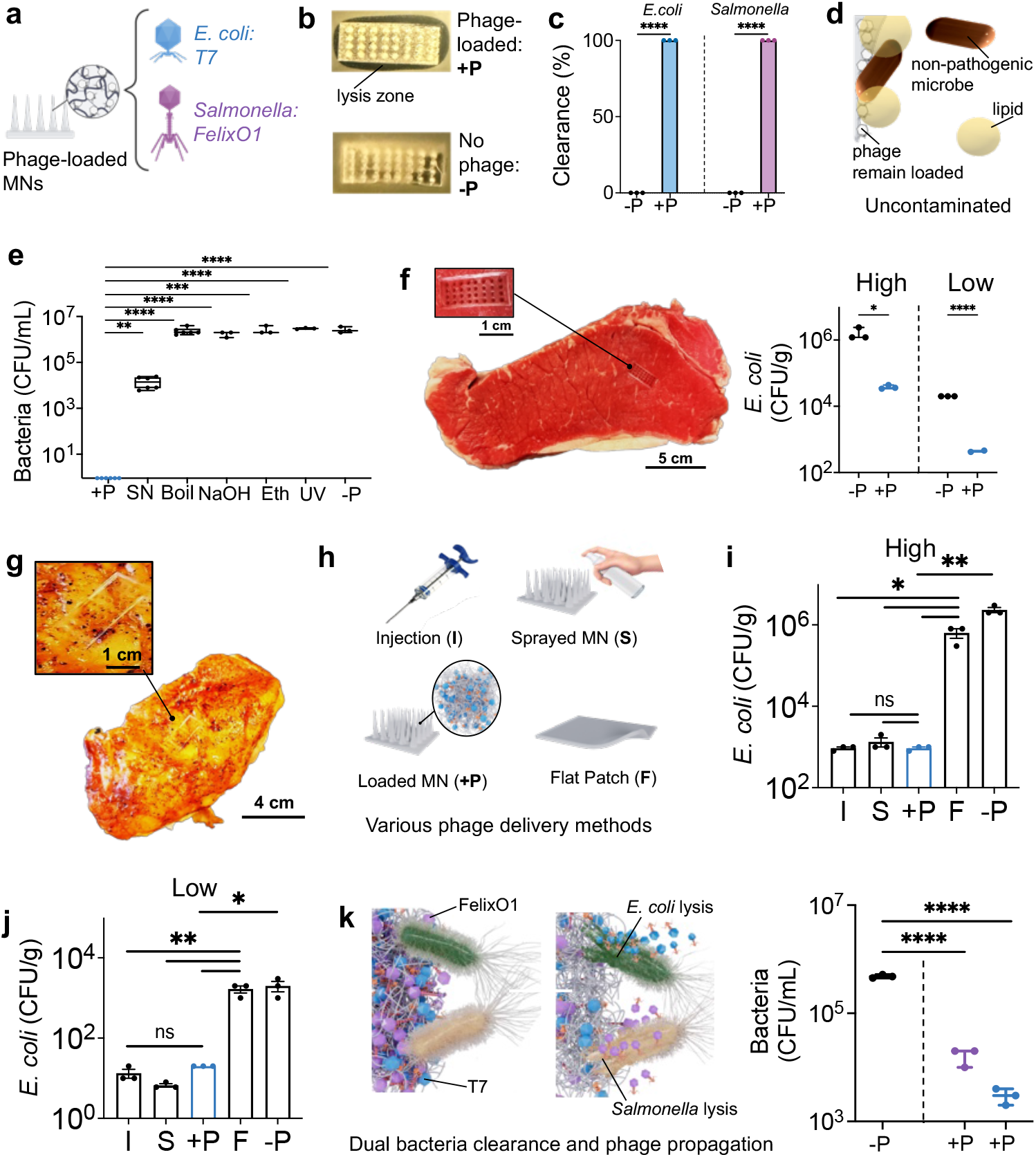
Full-system assessment of bacteriophage-loaded PMMA microneedles. (a) T7/FelixO1 bacteriophage-loaded microneedles. (b) Bacterial lawns with lysis zone shown for bacteriophage-loaded microneedles (+P) and absence of lysis zone with unloaded microneedles (−P). (c) Baseline *E. coli* and *Salmonella* decontamination with loaded microneedle arrays. (d) Bacteriophage remain loaded in the presence of other food constituents including lipid molecules and non-pathogenic microbes to prevent nonspecific delivery. (e) Bacteriophage degradation when arrays were subjected to sonication (SN), boiling water (Boil), sodium hydroxide (NaOH), 70% ethanol (Eth), and 1 hour ultraviolet light exposure (UV). (f) *E. coli* decontamination in steak with high (10^5^ CFU/g, left) and low (10^2^ CFU/g, right) levels of initial contamination. (g) Bacteriophage-loaded microneedles embedded in RTE chicken. (h) Various bacteriophage delivery methods including more invasive injection (I), non-specific sprayed microneedle arrays (S), and surface-only flat bacteriophage-loaded patches (F). (i-j) Performance assessment of various bacteriophage delivery methods within RTE chicken with high (i) and low (j) levels of initial *E. coli* contamination. (k) Dual-pathogen clearance with multiplex bacteriophage-loaded microneedles containing both *E. coli* and *Salmonella*-specific bacteriophage. All reported values represent the mean of all samples with error bars representing standard error of the mean. All asterisks represent significance differences at corresponding significance levels. (a) and (d) created using BioRender.

Based on the high prevalence of *E. coli* contamination in raw beef, food testing was first briefly performed on steak with high (10^5^ CFU/g) and low (10^2^ CFU/g) levels of contamination (**Figure 4f**). In both cases, samples with bacteriophage-loaded microneedles demonstrated significantly lower contamination, achieving nearly 2-logs reductions (*P* < 0.05, 0.0001). However, given the higher risk associated with RTE products, subsequent testing was performed with cooked chicken (**Figure 4g**). RTE chicken was tested with bacteriophage-loaded microneedles and compared to alternative delivery methods such as direct bacteriophage injection, bacteriophage-sprayed microneedles, and surface-level flat bacteriophage-loaded patches (**Figure 4h**). Again, both high (**Figure 4i**) and low bacterial (**Figure 4j**) contamination was induced, followed by an 18 hour incubation. In both cases, the bacteriophage-loaded patches did not exhibit significant differences in bacterial reduction compared to the more the direct injection and sprayed-microneedle approaches, while concurrently outperforming the flat bacteriophage-loaded patches. Specifically, chicken samples treated with the phage-loaded microneedles showed contamination levels of 10^2^ and 10^1^ CFU/g compared to 10^6^ and 10^3^ CFU/g in samples treated with unloaded microneedles, for both high and low levels of initial contamination, respectively. Thus, the phage-loaded microneedles demonstrated log-reductions of 3.40 and 2.00, compared to 0.57 and 0.08 for the flat patches (*P* < 0.05, 0.01). Importantly, bacterial reduction met regulatory thresholds of 10^1^ CFU/g in samples with low levels of contamination, effectively decontaminating the chicken.

To ensure widespread applicability, we combined both T7 and FelixO1 bacteriophage for dual-pathogen decontamination to create a multiplex antibacterial platform (**Figure 4k**). This system demonstrated significant contamination reductions of 1.46 and 2.20 logs for *Salmonella* and *E. coli*, respectively (*P* < 0.0001, 0.0001). Finally, the feasibility of a large-scale system was also explored by combining multiple small microneedle patches to create a unified, larger structure (**Method S3, Figure S10**). Such a system has the potential for decontaminating entire food products while integrating this technology into consumer applications, ensuring pathogen-specific safety throughout the food production pipeline.

## Conclusion

We have developed food-safe, bacteriophage-loaded microneedles, which enable antibacterial delivery directly into food, high-risk, fluid-rich matrices – all without any disruptions to organoleptic properties. This work enables the cost-efficient and effective decontamination of a myriad of food products from multiple bacterial contaminants. To ensure widespread applicability to a diverse range of food products, we first compared the performance of various polymeric materials through mechanical and stability characterization as well as through food-specific validation studies to ultimately determine PMMA as the ideal candidate material. While microneedle arrays pose immense promise within the food monitoring industry, the lack of research towards this application has limited their use in the space. Such food-centric testing represents the only extensive characterization of microneedles for food-based application and stands to provide a foundation for future works. Extensive performance assessment using contaminated whole chicken and meat products was then performed, demonstrating an over three-log improvement over surface-treatment approaches, comparable to the reduction observed through disruptive bacteriophage injection. Under some test conditions, the degree of pathogen reduction was sufficient to render the product safe to consume, in accordance with established regulatory limits. The widespread applicability of this system was also validated through multiplex antibacterial activity, where microneedle patches were loaded with two bacteriophage simultaneously. The developed platform can revolutionize current food contamination practices, preventing foodborne illness and waste through the active decontamination of food products. Its customizability, food-safe nature, and cost-effectiveness makes it well-suited for applications within the food industry.

## Methods

### Mold fabrication

All microneedle arrays were created using micromolding techniques. The positive microneedle resin molds and their associated 2D drawings were created using 3D computer assisted design (CAD) software (Autodesk Fusion) and then 3D printed using stereolithography (SLA) (B9Creations). The resin molds were smoothening with 99% isopropyl to ensure high resolution without the presence of artifacts. PDMS Sylgard 184 (Dow Inc.) negative molds were then casted to form master molds using a 10:1 ratio of base resin to curing agent. The casted molds were desiccated for 1 hour and then heat-cured at 60^°^C for 18 hours.

### Microneedle fabrication

PMMA microneedles were created by dissolving PMMA powder (MilliporeSigma) with 30% w/v of ethyl lactate or ethyl pyruvate solvents. This solution was thoroughly vortexed and then casted into the PDMS molds. Following a one-hour desiccation, the arrays were heat cured at 37^°^C overnight and then separated out of the molds. PVA microneedles were developed by creating a 12% solution of PVA powder (MilliporeSigma) and water. This solution was then casted onto heated PDMS molds and placed under vacuum at 80^°^C for 5 minutes and then placed at –20°C for 24 h. To create PDMS microneedles, PDMS was mixed in a 10:1 ratio of base to curing agent. The PDMS negative molds were then briefly treated with a 10% surfactant to avoid the microneedles from fusing to the PDMS mold. Once treated, the PDMS microneedles were casted in a similar desiccation and overnight heat treatment approach. Gelatin microneedles were created using a 15% w/v gelatin solution. This solution was stirred, bloomed for 5 minutes using a microwave and casted onto PDMS molds. The molds then underwent a heated vacuum treatment followed by a –20°C incubation in a similar manner as the PVA microneedles. For all microneedle materials, any visible air pockets post-vacuum desiccation were removed using a sharp needle.

### Microneedle imaging

Optical images of the microneedles were obtained using an iPhone 14 Pro smartphone. Fluorescence imaging was performed with an inverted microscope (NikonEclipse Ti2, Nikon Instruments Inc.) and was used to quantify fluorescence background. All material candidates were imaged across the DAPI, FITC, TRITC, and Cy5 fluorescence wavelengths at magnifications of 4X and 10X. Pixel values were quantified and averaged to determine background fluorescence levels.

### Scanning electron microscopy

Scanning electron microscopy images were obtained by isolating individual microneedles and sputter coating (Polaron E1500, Polaron Equipment Ltd., Watford, Hertfordshire) them with 5 nm of platinum. Samples were imaged using the JEOL JSM-7000F.

### Microneedle mechanical testing

Individual microneedles were isolated for compressive testing, with a minimum of triplicate samples for each material at each force. A micromechanical tester (Biomomentum Mach 1) then used to administer forces of 50, 100, 250, and 500 gf at a velocity of 0.75 mm/s. Each test was recorded with images acquired at a sampling rate of 5 seconds. Height reductions were then quantified using image analysis software (ImageJ). Stability assessments for the storage, pH, and sonication tests was performed with halved microneedle patches that were precisely cut. During testing, any needles that were damaged and broken as a result of the applied force were indicated as not reusable. For each agarose concentration, triplicate samples were used to penetrate the disks, and the number of holes penetrated were recorded.

### Microneedle penetration and reusability in food

All five food products were prepared in approximately 2 cm x 2 cm samples. Halved microneedle patches were then applied to the top surface of each food product and then constant penetration forces 250, 500 and 750 gf were applied at a velocity of 0.75 mm/s. Penetration was then recorded as the total number of penetration holes as a percentage of the total needles within the patch. Reusability was quantified as a percentage of the number of undamaged needles that remained within the microneedle patch compared to the total number of needles on the patch.

### Microneedle penetration in food-industry procedures and in food matrix fluids

Microneedle patches were inserted into 5 cm x 5 cm pieces of the respective food products using a constant force of 500 gf at 0.75 mm/s using the mechanical tester. Tumbling tests were performed with microneedle patches inserted into peaches and mushrooms. These samples were then subjected to tumbling at 500 rpm for 2 hours. Vacuum tests were performed with microneedles inserted into cheese pieces. The samples were placed under and removed from vacuum repeatedly over five cycles. Any microneedle patches that were retracted from the cheese between cycles were removed from further cycles. Chicken purge and saline solution testing involved first inserting microneedles into chicken and fish pieces, respectively, and then applying 1 mL of the respective solution onto the food item. These samples were incubated for 24 hours to ensure sufficient time for any interactions between the solutions and microneedle materials. Following all tests, the total holes from penetration and any instances of broken or deformed microneedles were documented.

### T7 phage propagation

T7 phage was propagated using *Escherichia coli* strain K12 BL21 (Sigma-Aldrich, CMC001) as a host. A 5 mL pre-culture of *E. coli* was prepared in Luria-Bertani (LB)-Miller broth in a shaking incubator set to 180 rpm and 37 °C overnight. The following day, a fresh 5 mL 1:200 subculture was made using LB-Miller broth and grown to an optical density at 600 nm (OD_600nm_) of 0.6. 10 µL of T7 phage (~ 10^8^ PFU/mL) was then added to the subculture followed by 50 µL of 1M CaCl_2_ solution to assist in phage infectivity. The T7-subculture solution was then incubated in a shaking incubator set at 180 rpm and 37 °C for 8 hours. The lysate was centrifuged at 7000×g for 15 minutes and the bacteria pellet was discarded. The phage-containing supernatant was filtered through a 0.2 µm filter (Fisher Scientific, 13100106) to remove residual bacteria. 5 mL of polyethylene glycol (PEG) was added to the filtered supernatant and stored overnight at 4 °C. The following day, the PEG-phage solution was centrifuged at 5000×g for 60 minutes. The resulting phage pellet was resuspended in sterile RO water.

### FelixO1 phage propagation

*Salmonella enterica* serovar Typhimurium (ATCC 700720). An overnight culture of *S. enterica* was prepared by inoculation of 3 mL of Tryptic Soy Broth (TSB) from a frozen glycerol stock followed by incubation at 180 rpm and 37 °C for 16-18 hours. For propagation of the phage, 100 µL of overnight *S. enterica* culture and 100 µL of FelixO1 stock (10^5^ PFU/mL) were added to 30 mL of TSB and incubated at 180 rpm and 37 °C for approximately 24 hours. Following the propagation, 10 µL of chloroform was added to the suspension and allowed to incubate at room temperature for 15 minutes. Crude lysate was then centrifuged at 7000xg for 15 minutes and the bacteria debris discarded. The supernatant containing FelixO1 was collected, sterilized by filtration using 0.2 µm filter (Fisher Scientific, 13100106), and stored at 4 °C.

### Bacteria culture

*E. coli* K12 and *Salmonella enterica* were cultured in LB and TSB media, respectively, for 18 hours at 37 °C under constant agitation at 180 rpm from glycerol stock solutions. This solution was used directly for antibacterial bacterial lawns. For contamination experiments, the overnight bacteria stock was centrifuged at 7000 rpm for 15 minutes and then resuspended in phosphate-buffered saline (PBS) buffer.

### Bacteriophage-incorporation into microneedles

PMMA solution was created as outlined in *Microneedle fabrication*. Once the solution was vortexed, 25% v/v bacteriophage was added. The bacteriophage-loaded solution was vortexed at a low speed to avoid unnecessary bacteriophage damage and then the resultant solution was casted in the same approach as previously outlined. For the development of the multiplexed platform, both bacteriophage were added to the PMMA solution in equal parts and casted in the same manner.

### Antibacterial testing

Antibacterial testing for the bacteriophage-loaded microneedles was performed using bacterial lawns. Overnight bacterial cultures were produced as outlined in the *Bacterial Preparation*. LB agar plates were produced by suspending LB powder (Fisher Scientific) and 1.5% w/v agar in sterile water, autoclaving the solution and dispensing into sterile petri dishes (Fisher Scientific). Soft agar overlays were created by suspending LB powder and 0.6% w/v agar in sterile water and aliquoting this solution into glass test tubes. Bacterial lawns were prepared by adding 150 μL of bacteria within 3 mL of soft agar, and this solution was vortexed and then poured onto LB agar plates. Both loaded and unloaded microneedle arrays were inserted into the top layer of the bacterial lawn and incubated overnight within a stationary incubator at 37°C. Baseline bacterial clearance in solution was assessed by serial dilution on MacConkey agar plates (SigmaAldrich) for both bacteria using spot plating techniques. Plaque assays were also similarly performed on bacterial lawns with spot plating to quantify bacteriophage delivery. Bacteriophage suspensions were added into solutions of PBS, peach pulp, and chicken purge, serially diluted, and then plated.

### Phage degradation study

Three bacteriophage-loaded microneedle patches were used as samples for each stressor to provide triplicate measurements for each condition. Samples which underwent sonication were placed in 50 mL of water and sonicated for 1 hour. Samples exposed to ultraviolet light were placed under the germicidal UV-C light of a biosafety cabinet for 1 hour. All other samples were submerged in 50 mL of either boiling water for one hour or within 1 M sodium hydroxide or 70% ethanol overnight. All samples submerged in sodium hydroxide and ethanol were thoroughly rinsed in water to avoid nonspecific bacterial reductions from the harsh chemicals. Finally, all samples were incubated in 1 mL of 10^6^ CFU/mL of *E. coli* for 18 hours at 37 °C along with positive and negative controls of intact loaded microneedles and unloaded microneedles, respectively. Following this incubation, all samples were vortexed, serially diluted, and selectively plated. The plates were placed in a stationary incubator at 37 °C for an overnight incubation, after which bacterial growth was quantified.

### Antibacterial assessment within contaminated food products

To avoid confounding growth from nonspecific microbes innately present within the food matrix, all food testing was performed with selective MacConkey agar plates. Both raw beef and chicken pieces were prepared into 200 g sized pieces and injected with a volume of *E. coli* equivalent to 10^5^ and 10^2^ CFU/g of contamination. Importantly, contamination was induced about 2-3 cm below the surface and in multiple locations throughout the food sample to best emulate contamination. Three bacteriophage-loaded microneedle patches and unloaded controls were then applied to each sample and then the samples were incubated for 18 hours. Raw steak samples were incubated at 4 °C while RTE chicken was incubated at 37 °C in accordance with their standard storage temperatures. Post incubation, the food products were processed in a stomaching approach to extract sample solution. This solution was serially diluted and selectively plated in the same approach as the *Phage degradation study*.

### Phage injection, spraying and loading into flat patches

Bacteriophage injection consisted of directly injecting 1 mL of bacteriophage in one location about 1 cm below the chicken surface. Sprayed microneedles were created by spraying 1 mL of bacteriophage directly on an unloaded microneedle array using a small atomizer. Finally, flat bacteriophage-loaded patches were developed using the same approach the bacteriophage-loaded microneedles, except using a custom flat CAD mold that was created without the presence of any needles.

## Supporting Information

Supporting Information is available online.

## Acknowledgements

A.P. is a recipient of the Vanier Canada Graduate Scholarship and S.K. is a recipient of the Banting Fellowship, both awarded by the Natural Sciences and Engineering Research Council of Canada. This work was supported by the Ontario Early Researcher Award, Mitacs and Toyota Tsusho Canada Inc. grants to T.F.D. T.F.D. and Z.H. acknowledge funding from the NSERC Discovery Grants, Canada Research Chairs, and Ontario Early Researcher Award Programs.

