## Supplementary Information for "Bacteriophage-Loaded Microneedle Patches for Targeted and Minimally Disruptive Foodborne Pathogen Decontamination"

**Note S1: Choice of microneedle polymers**

Microneedle research has surged over the past few years, with thousands of papers published in literature exploring various fabrication methods and materials.[1,2] Polymeric materials, including silicone, have emerged as the optimal material choice for microneedles given their low cost and tunable mechanical properties.[3] Other materials such as metals were excluded from this work given their high cost and requirement for more specialized manufacturing procedures.[4] While the selected polymeric candidate materials were molded to create solid microneedles, they can also be used to create dissolvable, swellable and hollow microneedles through various manufacturing techniques.[4] The selected polymers all had low to moderate price points and could be easily fabricated through micromolding techniques in a cost efficient manner. Importantly they were all biocompatible and either biodegradable or recyclable.[3] Some properties of these materials are summarized in **Supplementary Table 1** and their polymeric reactions are described in **Supplementary Figure 1**.

**Note S2: Selection of bacteriophages**

The T7 phage used in this work is highly specific to *E. coli* ER2738 as well as *E. coli* BL21 and belongs to the *Podoviridae* or short-tailed family. The FelixO1 phage offers species level specificity to *Salmonella*, belonging to the *Myoviridae* or long-tailed family. While both phages offer high antimicrobial activity, it is important to note that various mixtures of different phages must be used for polymicrobial decontamination due to the high specificity that each phage offers.[5] An important consideration for the development of this platform was ensuring that the selected phages could withstand the micromolding process and maintain their infectivity. In food specific applications, T7 has shown relatively high stability, especially in highly acidic environments, making it a strong candidate for use in the final system.[6] While the stability of FelixO1 is less characterized, *Myoviridae* phage have shown sufficient stability in various organic solvents, indicative of their compatibility with the developed system.[7] The use of lytic phage was also a key consideration as some bacteriophage, such as the well-characterized M13 phage, follow lysogenic reproduction cycles which do not kill their host bacteria and thus cannot provide any decontamination.

**Method S1:** *Microneedle permeability assessment*

Biomolecule permeability was assessed by loading a pH-responsive, food-derived, and food-safe anthocyanin dye within the polymer solution prior to casting. The microneedles were cured through the same process as before and then peeled from their molds. These patches were carefully imaged using a scanner to standardize the imaging process. The anthocyanin-loaded microneedles were then placed in 5 mL of 1 M sodium hydroxide (NaOH) solution for 10 mins, 3 days, and 6 days. Following this incubation, the microneedle patches were carefully removed and dried. The patches were reimaged using the same conditions with the scanner. Post imaging, the colour change of the pre-incubation and post-incubation was quantified using ImageJ. Specifically, the average pixel value for the entire patch was recorded with triplicate readings across three different patches. This was repeated for all four polymer materials. Given that the red cabbage anthocyanin used for this study changes colour from purple to green in strong basic environments, all colour change was assessed using the green pixel values from the RGB images. Anthocyanin dye dissolved in an aqueous solution was used as positive controls, where the colour change displayed in solution represented 100% permeability. The color change for all the polymeric materials was then normalized against this permeability to quantify the overall permeability of the microneedle array. These results are summarized in **Supplementary Figure 6**.

**Method S2:** *Porcine skin penetration testing*

Porcine skin was sliced into approximately 2 cm x 2 cm samples and placed on the base of the micromechanical tester. Halved microneedle patches were placed on top of the porcine skin sample and a constant force was applied to the patch at a velocity of 0.75 mm/s. Forces of 48 N and 72 N were both used and were normalized to 3 N/needle and 4.5 N/needle for each 16-

needle patch in accordance with other reported studies in literature. Penetration was then recorded as the total number of penetration holes as a percentage of the total needles within the patch. Reusability was quantified as a percentage of the number of undamaged needles that remained within the microneedle patch compared to the total number of needles on the patch. These results are summarized in **Supplementary Figure 7**.

**Method S3:** *Development of a large-scale microneedle array*

A base was first created with 3D computer assisted design (CAD) software (Autodesk Fusion). The base was then 3D printed with a PLA filament printer (Ender 3 V2, Shenzhen Creality 3D Technology Co., Ltd., China). This base was built according to the dimensions of a standard household food storage container (**Figure 1f**) but can be customized to any size as 3D printing is a highly tunable manufacturing method. An adhesive backing was then applied to the base. Individual PMMA microneedle patches, each consisting of 32 individual microneedles, were applied onto the base along 8 rows and 7 columns. In total, the complete unified structure consists of 56 microneedle patches with 1792 microneedles (**Supplementary Figure 10**). Such a system can be used to offer whole-product, continuous decontamination without significant effects of the organoleptic properties of the food product.

**Table S1: Overview of microneedle material properties**

| Material | Tensile Strength (MPa) [8,9] | Cost[8] | FDA GRAS | Recyclable | Bio-degradable | Types of Micro-needles | Key Details[1,8] |
|---|---|---|---|---|---|---|---|
| PMMA | 62 | $$ | ✓ | ✓ | X | Solid, Hollow | Easy, high resolution fabrication |
| PVA | 48 | $ | ✓ | ✓ | ✓ | Solid, Dissolvable | Good hydrogel micro-needles when paired with other polymers |
| PDMS | 25 | $$ | ✓ | ✓ | X | Solid, Dissolvable | More often used for casting micro-needle molds |
| Gelatin | 0.021 | $ | ✓ | ✓ | ✓ | Solid, Swellable | Tunable strength when paired with other materials |

$ < $25/kg    $$ < $100/kg    $$$ > $100/kg

**Table S2: Summary of microneedle material mechanical characterization**

| Material | Reusability at 10 N Force (%) | | Reusability after 4 weeks (%) | | Reusability after storage in an acidic (pH = 6) environment (%) | | Reusability after 10 mins of sonication (%) | | Penetration of a 1.5% (w/v) agarose disk (%) | | Water absorption after 1h (wt% increase) | |
|---|---|---|---|---|---|---|---|---|---|---|---|---|
| PMMA | 91 | ↑↑↑ | 66 | ↑↑ | 38 | ↑ | 48 | ↑↑ | 100 | ↑↑↑ | 1 | – |
| PVA | 90 | ↑↑↑ | 18 | ↑ | 0 | – | 71 | ↑↑ | 100 | ↑↑↑ | 0 | – |
| PDMS | 100 | ↑↑↑ | 100 | ↑↑↑ | 100 | ↑↑↑ | 98 | ↑↑↑ | 20 | ↑ | 31 | ↑ |
| Gelatin | 19 | ↑ | 0 | – | 0 | – | 0 | – | 0 | – | 125 | ↑↑↑ |

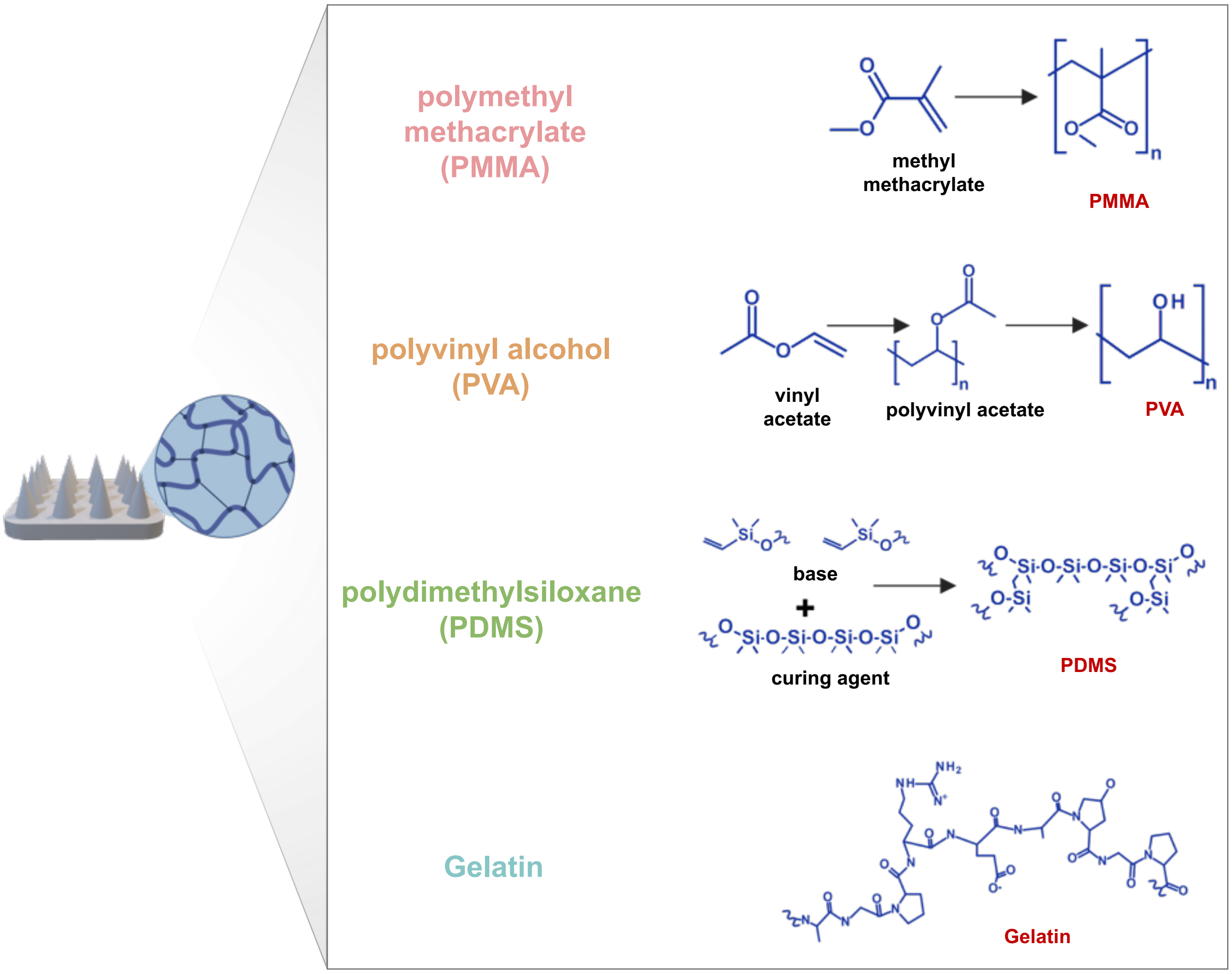


**Figure S1.** Chemical compounds of biocompatible polymers used for microneedle fabrication labelled in red along with relevant monomers/bases labelled in black. While PMMA and PVA are formed using free radical polymerization, commercially available powders were used for both in this work. PDMS is created by mixing the prepolymer base agent with a curing agent in a 10:1 weight ratio after undergoing various intermediary reactions. As a naturally occurring polymer, gelatin does not have any polymerization reactions. Commercially available gelatin powder was used for this work.

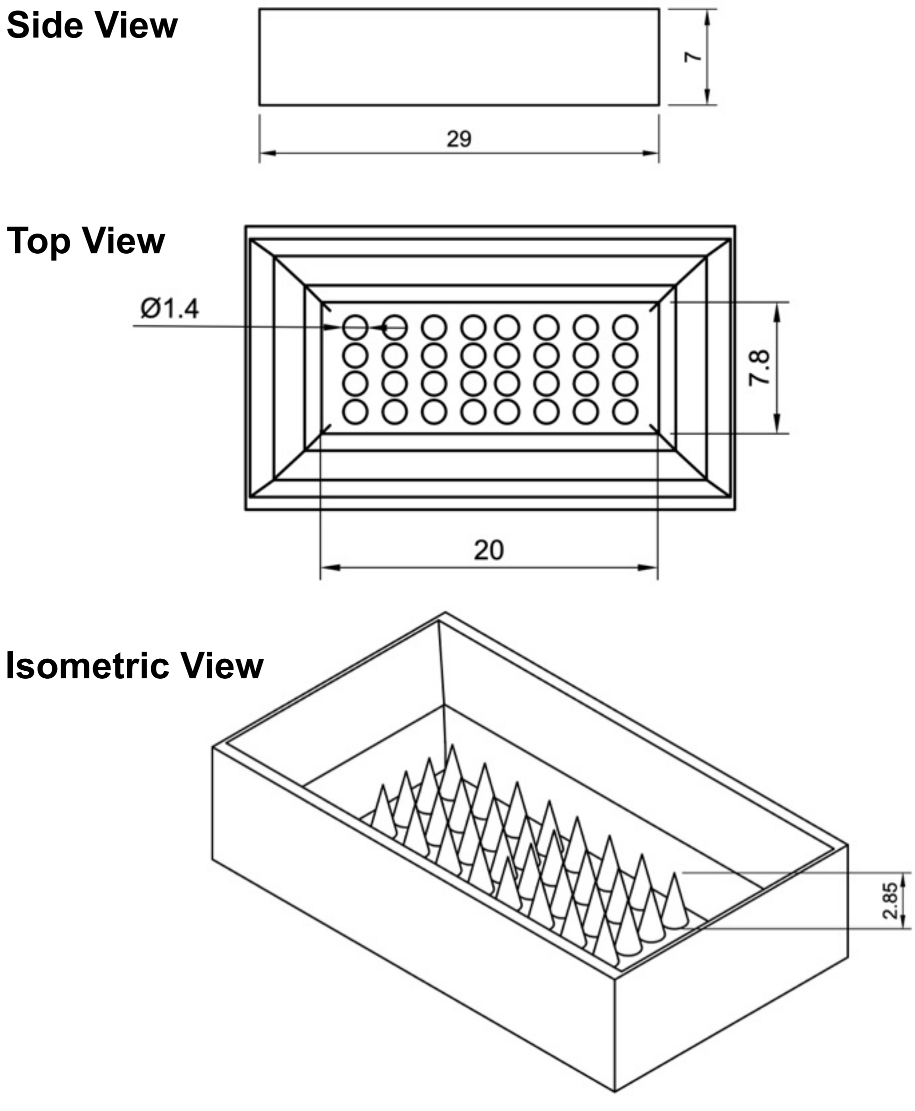


**Figure S2.** 2D drawings of 3D-modelled microneedle molds with pertinent dimensions shown in millimetres for side, top, and isometric views. Microneedle diameter and height highlighted in top and isometric views, respectively.

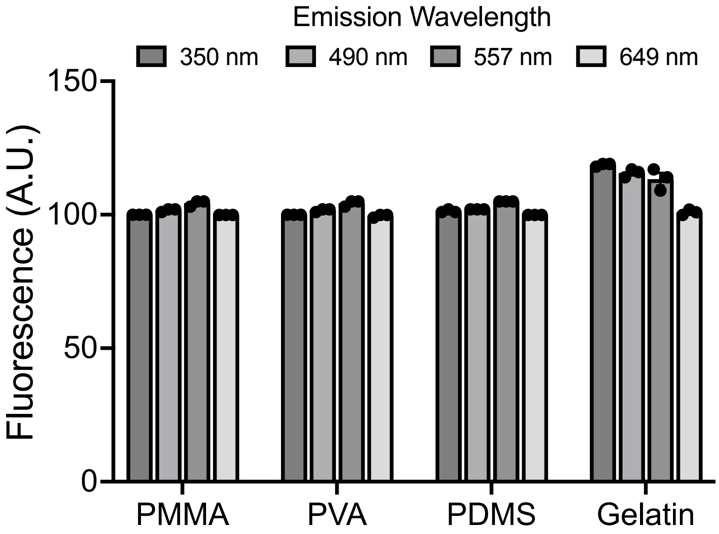


**Figure S3.** Mean background fluorescence of selected polymer candidate materials. Reported values represent the mean of triplicate samples with error bars representing sample standard deviation. The reported emission wavelength channels roughly correspond with the commonly used diamidino-2-phenylindole (DAPI), fluorescein isothiocyanate (FITC), tetramethylrhodamine (TRITC), and Cy5 fluorescence dyes. These dyes can be used to achieve blue, green, orange, and red fluorescence microneedle arrays, respectively, particularly for biosensing applications.[10]

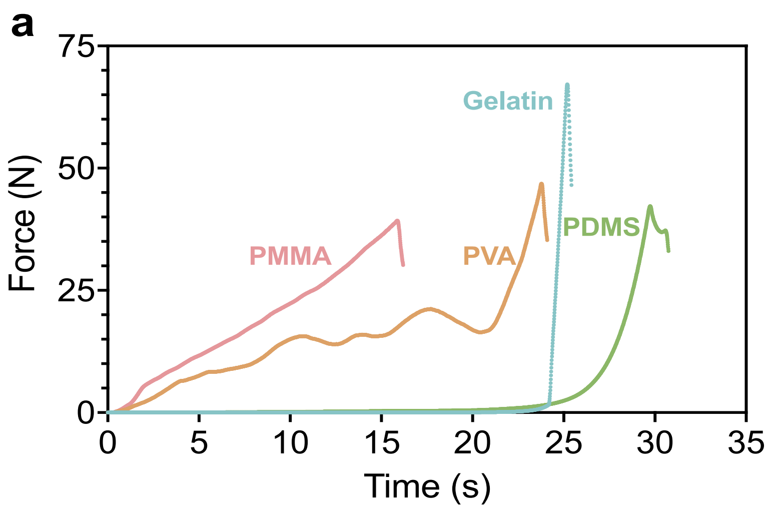


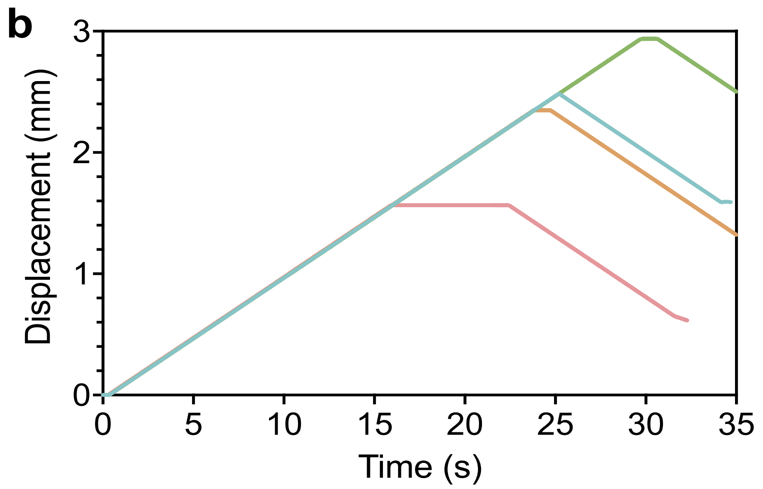


**Figure S4.** Force versus time (a) and displacement versus time (b) graphs of all material candidates under 50N of compressive force. For (a), the end of each curve signifies material fracture or complete compression, with plateaus in (b) representing corresponding displacement. For (b), a positive change in displacement corresponds to downward motion as needle compresses. A maximum displacement of 3 mm in (b) was used given that the maximum height of each microneedle is 2.85 mm.

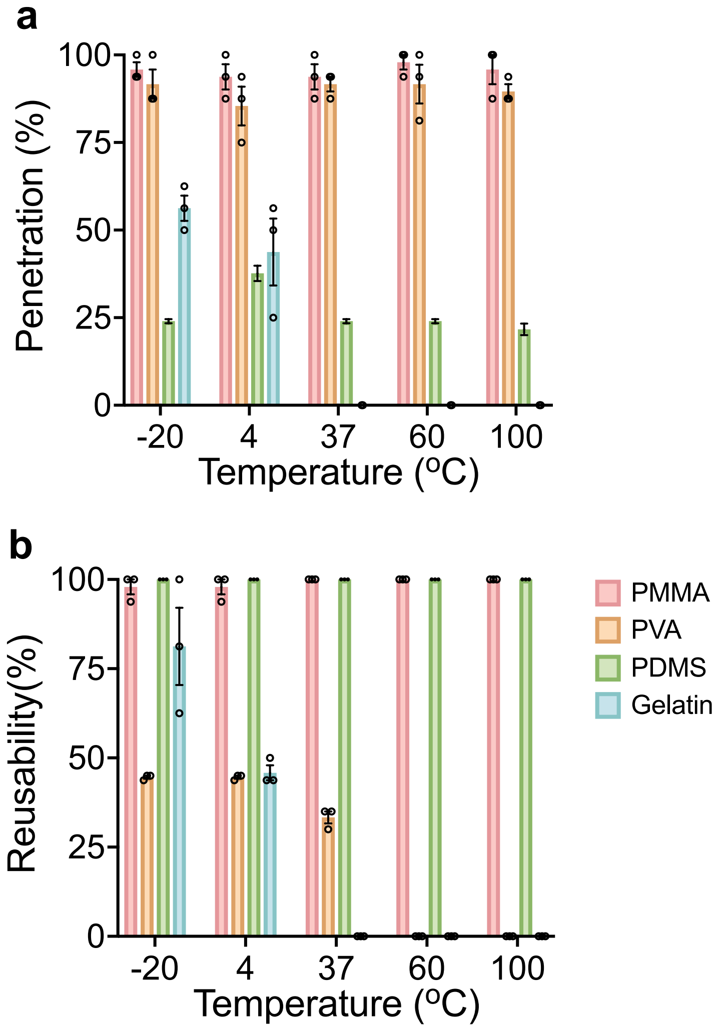


**Figure S5.** Microneedle stability assessment based on penetration (a) and reusability (b) evaluation after storage at various temperatures ranging from -20 to 100 °C. Reported values represent the mean of triplicate samples with error bars representing standard error of the mean.

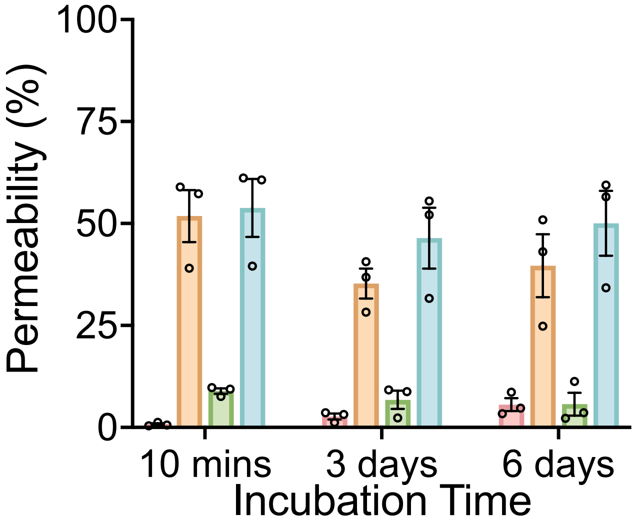


**Figure S6.** Permeability of selected polymer candidate materials based on change in colour of a pH-sensitive dye when dissolved in reference solution. Incubation time corresponds to length of storage within sodium hydroxide (NaOH) to illicit a colour change. Reported values represent the mean of triplicate samples with error bars representing standard error of the mean.

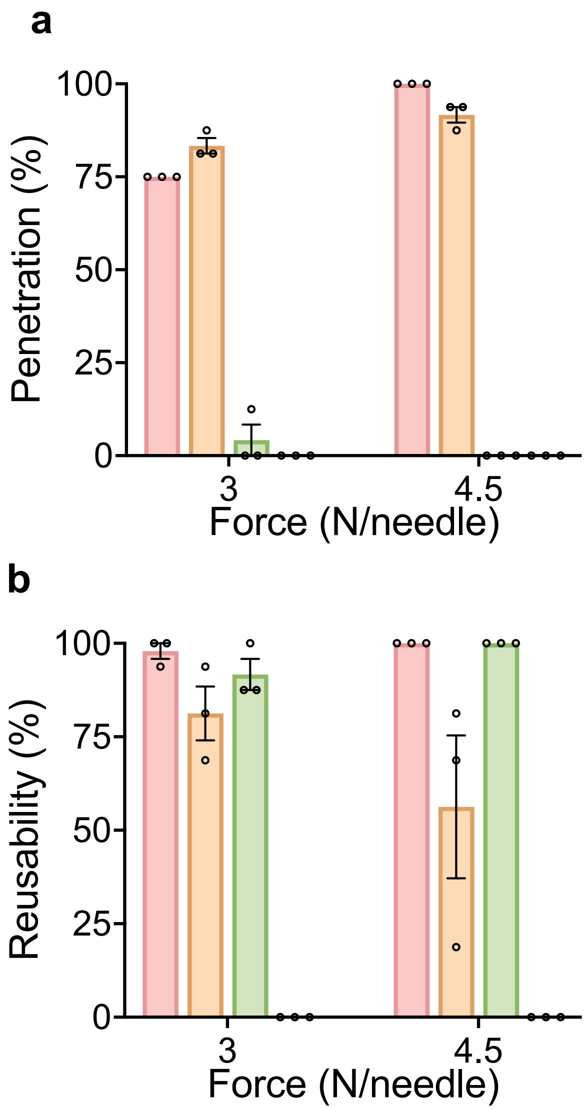


**Figure S7.** Penetration (a) and reusability (b) of microneedles when inserted into porcine skin at forces of 3 and 4.5 N/needle. Reported values represent the mean of triplicate samples with error bars representing standard error of the mean.

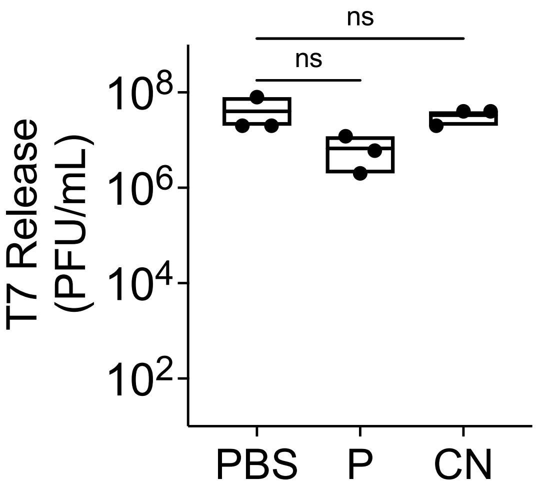


**Figure S8.** Bacteriophage release with peach fluid and chicken purge compared to buffer controls. Reported values represent the mean of triplicate samples with error bars representing standard error of the mean.

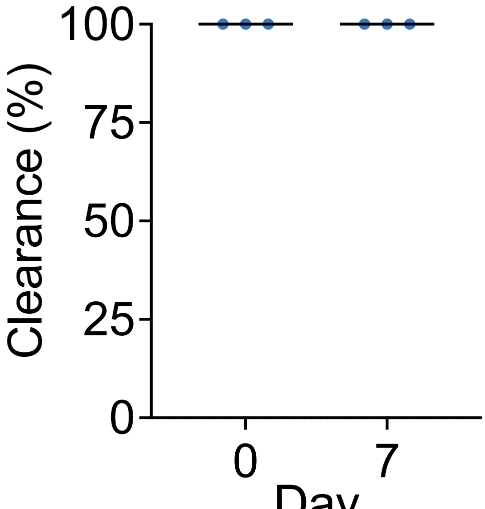


**Figure S9.** Bacteriophage delivery with peach fluid and chicken purge compared to buffer controls. Reported values represent the mean of triplicate samples with error bars representing standard error of the mean.

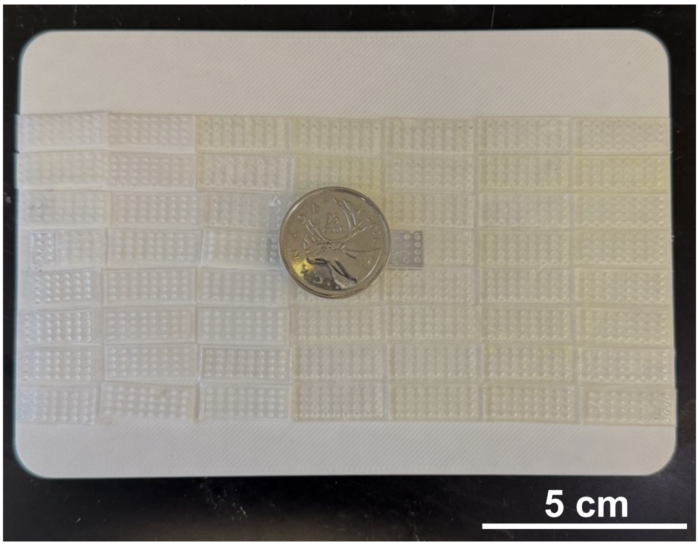


**Figure S10.** Potential large-scale use of numerous microneedle arrays for whole-product, continuous decontamination. Coin used to show relative scale.
